# Causal Evidence for the Dependence of the Magnitude Effect on Dorsolateral Prefrontal Cortex

**DOI:** 10.1101/306746

**Authors:** Ian C. Ballard, Gökhan Aydogan, Bokyung Kim, Samuel M. McClure

## Abstract

Impulsivity refers to the tendency to insufficiently consider alternatives or to overvalue rewards that are available immediately. Impulsivity is a hallmark of human decision making with well documented health and financial ramifications. Numerous contextual changes and framing manipulations powerfully influence impulsivity. One of the most robust such phenomenon is the finding that people are more patient as the values of choice options are increased. This magnitude effect has been related to cognitive control mechanisms in the dorsal lateral prefrontal cortex (dlPFC). We used repetitive transcranial magnetic stimulation (rTMS) to transiently disrupt dlPFC neural activity. This manipulation dramatically reduced the magnitude effect, establishing causal evidence that the magnitude effect depends on dlPFC.

## 1 Introduction

Humans are often short-sighted in their decision-making and choose immediate gratification at the expense of great longer-term benefits. This behavior is pervasive and contributes to large scale societal problems such as obesity^1^, drug addiction ^2^, and retirement^3^ and health savings shortfalls^4^. A prevalent laboratory measure of impulsivity is the discount rate estimated from intertemporal choice tasks. Individual discount rates are stable across time^5^, but vary substantially based on choice context and framing^6^. One of the largest contextual effects known to reduce impulsivity is the *magnitude effect*, which refers to the phenomenon that people have lower discount rates when choosing between high magnitude options compared to choices between lower valued rewards^7^. Recent behavioral studies support the idea that large magnitudes trigger increased self-control, which in turn reduces impulsivity^8^. Further, correlational fMRI investigations have associated high magnitude decision-making with activation of the dorsolateral prefrontal cortex (dlPFC)^8^. We tested whether we could find causal evidence that dlPFC-mediated control underlies the reduced impulsivity observed in the magnitude effect. To do so, we used repetitive transcranial magnetic stimulation (rTMS) to temporarily disrupt dlPFC activity and test for effects on the size of the magnitude effect.

Findings from neuroscience indicate that there are at least two means by which discount rates may be influenced in terms of brain systems involved. Regarding the dlPFC, it is well known that this region is activated when self-control is exerted^9^ and is necessary for the deployment of self-control^10^. The dlPFC likely influences control of decision making via multiple mechanisms: the dlPFC supports the maintenance of goals and task information^11-13^, increases the fidelity of internal representations of task variables^14^, and exerts top-down control over affective systems^15^. Each of these processes may permit longer-term goals to bias processing in other brain areas so as to reduce impulsivity.

A second means by which brain activity may be altered to reduce discount rates relates to activity in brain reward areas – the ventral striatum and ventromedial prefrontal cortex (vmPFC). Neurons in the ventral striatum encode the learned values of different choice attributes^16^, and in human neuroimaging experiments, ventral striatum activation tracks the subjective value of rewards^17^,^18^. Information from these and other circuits is integrated in the vmPFC^19^, where different options are compared sequentially^20^. It is feasible that contextual changes may alter how reward prospects are evaluated so as to change impulsivity without altering dlPFC activity.

Recent research findings indicate that manipulations that alter decision making can act by altering responses in dlFPC or brain reward structures. We have proposed that large reward magnitudes trigger engagement of control processes in the dlPFC^8^, which results in more patient decision-making, whereas manipulations that directly affect the evaluation of rewards primarily impact activity in the striatum^21^. We tested whether we could detect causal support for these modes of influencing delay discounting using rTMS, which transiently disrupts neural activity^22^, over the dlPFC. Beyond establishing causal evidence in support of fMRI results, this finding would also help adjudicate between alternative psychological and economic models of the magnitude effect. In particular, prior behavioral experiments that have attempted to rule out alternate models of the magnitude effect have been unable to exclude the *utility account*^6^,^8^.

According to this account, subjective valuation itself is sensitive to magnitudes, such that the ratio between the utilities of $20 and $10 is perceived to be smaller than the ratio between the utilities of $2,000 and $1,000 (i.e. increasing elasticity with magnitude). Because of this, the larger reward in the high-magnitude condition is perceived as proportionately more valuable than the larger reward in the low-magnitude condition. rTMS over dlPFC has been shown to increase impulsive choice without altering subjective valuation of rewards, consistent with a role for the dlPFC in self-control^10^. This finding is also consistent with substantial evidence that the medial wall of the prefrontal cortex is associated with valuation^18^,^23^, and this area should not be directly affected by rTMS over dlPFC. Because rTMS over dlPFC should not directly affect brain circuits involved in valuation, the utility account predicts that rTMS should have a similar effect in high-versus-low magnitude conditions. In contrast, the self-control model of the magnitude effect predicts that rTMS should have a larger effect in the high-magnitude than the low magnitude condition.

## 2 Results

### 2.1 Magnitude Effect

We hypothesized that rTMS over dlPFC would disrupt the magnitude effect. We used two control measures of baseline discount rate for high and low magnitude conditions. The first was measured on a different day prior to the experiment (pre-experiment) and the second was measured during sham rTMS (Sham). We tested all effects versus both controls. This approach provides a stronger test of our manipulation because it confirms that the effect of rTMS is robust to day-to-day fluctuations in impulsivity. We constructed a separate mixed-effects ANOVA for each measure of baseline. Both models had hemisphere as a between-subject effect and rTMS level and framing/context condition as within-subject effects. We first tested whether we reproduced the magnitude effect and found a robust effect of reward magnitude on discount rate with respect to both pre-experiment baseline, *F*(1,25) = 172, *p* < .001, □*g*^2^= 0.39 and sham rTMS, *F*(1,25) = 43.8, *p* < .001, □*g*^2^= 0.27, Figure 1. This effect is similar in size to that observed in previous studies^8^.

**Figure 1.**
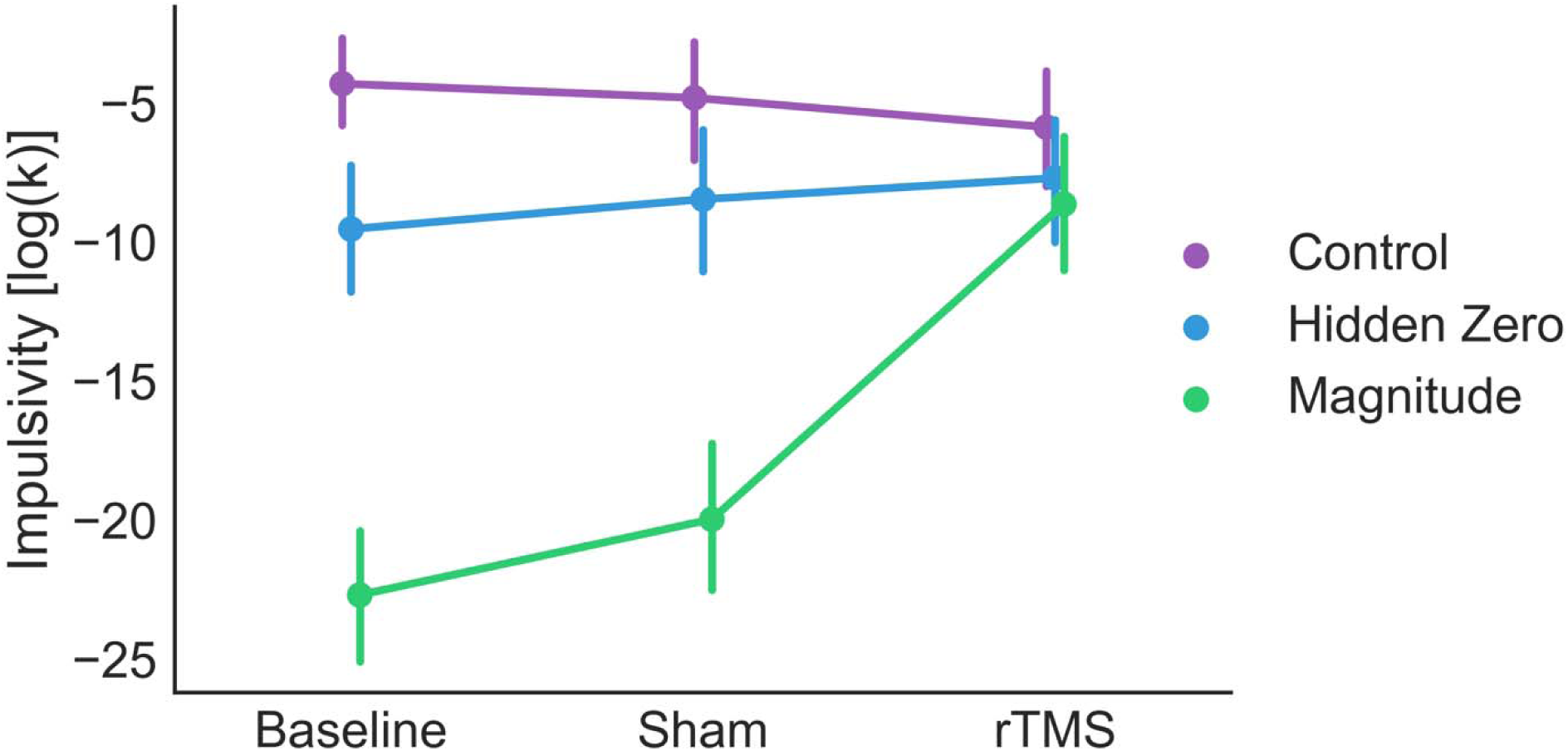
Impulsivity (discount rate, *k*) plotted as a function of rTMS condition and context/framing effect. Both the large magnitudes and explicit zero manipulations reduced impulsivity relative to control. rTMS reduced the magnitude effect relative to both baseline and sham. rTMS reduced the hidden zero effect relative to baseline but not to sham. Error bars reflect between-subjects standard errors of the mean.

Our central hypothesis was that rTMS over dlPFC should reduce the magnitude effect. We observed an interaction in which rTMS reduced the size of the magnitude effect with respect to both pre-experiment baseline, *F*(1,25) = 90.4, *p* < .001, □*g*^2^= 0.27, and sham rTMS, *F*(1,25) = 49.3, *p* < .001, □*g*^2^ = 0.15, Figure 2. Therefore, rTMS over dlPFC dramatically reduces the magnitude effect. Because significant interactions can complicate interpretation of main effects, we conducted several post-hoc *t*-tests to better understand these results. First, we tested whether we observed a significant magnitude effect in the absence of rTMS. Large magnitudes reduced impulsivity relative to both baseline, *t*(24) = 15.6, *p* < .001, and sham, *t*(24) = 7.9, *p* < .001, replicating the magnitude effect. We predicted that rTMS should have a larger impact on high magnitude choices. We examined whether rTMS increased impulsivity for the low magnitude choices and found no effect relative to either baseline in either hemisphere or for both hemispheres pooled together (all *p* > .15). Although this finding fails to replicate the finding that rTMS over dlPFC increases impulsivity^10^, an analysis using comparable statistical methods to previous work revealed a small effect of rTMS on low magnitude discount rates (Figure S1).

**Figure 2.**
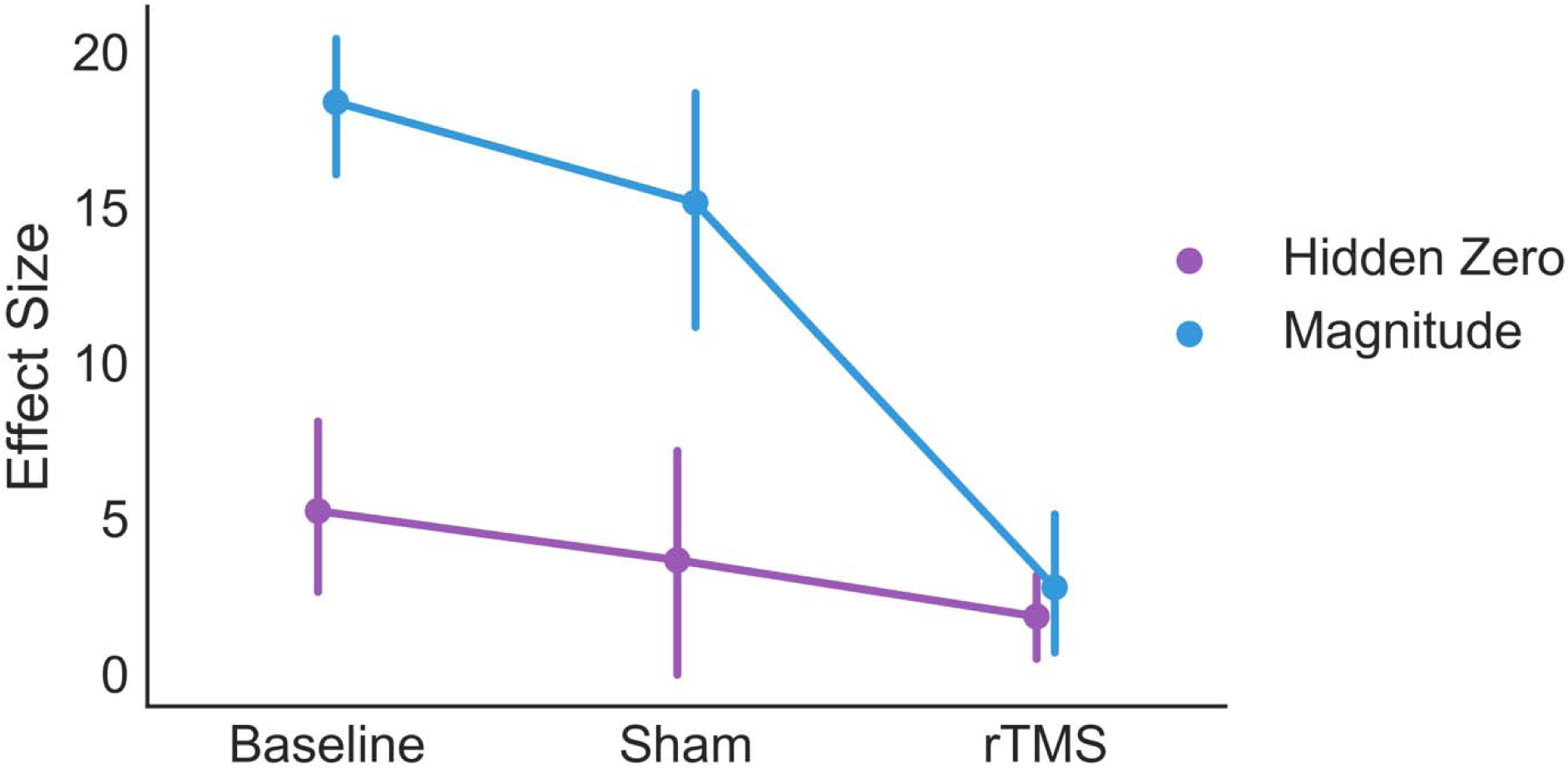
Comparison of the effect of TMS on the magnitude and hidden zero effects. “Effect size” refers to the difference in *log*(*k*) between the contextual/framing conditions and the control condition. The difference in impulsivity between conditions is plotted for each manipulation. Above zero effect size indicates an effect of choice framing or context. The change in effect size caused by TMS is larger for the magnitude effect than the hidden zero effect. Error bars reflect between-subjects standard errors of the mean.

We conducted control analyses to test whether these effects might arise from a general disruption of decision-making under rTMS. First, we examined whether rTMS might introduce noise into decision-making that eliminates the difference between manipulations. We did not find any effect of TMS on the decision noise parameter of the hyperbolic discounting model, *p* > .2. However, unlike discount rate, this parameter is poorly constrained and individual fits are noisy. We therefore conducted a more sensitive analysis in which we labeled choices as “correct” if the response was consistent with the discount rate estimated for that subject, and “incorrect” otherwise. We then tested whether rTMS influenced the number of correct choices, or conversely, the number of mistakes. If rTMS generally increased noise in responding, it should have increased the number of “incorrect” responses observed. Conversely, we find that rTMS actually increases the percentage of “correct” choices relative to both baseline, F(1,14) = 25.2, p < .001, and sham, F(1,14) = 20.3, p < .001, Figure S2.

Next, we conducted a related analysis to determine whether rTMS shifts individual discount rates without disrupting the relationship between subjective value and choice. Define *k*_*eq*_ to be the value of *k* that would result in a subject being indifferent between the two options in a given choice. If choices were ordered by *k*_*eq*_, a subject’s *k* specifies where in this ordered list they switch from preferring smaller-sooner to larger-later rewards. Consequently, the probability of choosing smaller-sooner rewards (*pSS*) should decrease monotonically with increasing *k*_*eq*_. If rTMS alters delay discounting by changing discount rate then it ought to change *pSS* while maintaining the relationship between *k*_*eq*_ and *pSS*. That is, a shift in discount rate will change the likelihood of choosing the smaller reward for all choices, but it will not alter the relative probability of choosing the smaller reward between choices. Alternatively, rTMS could disrupt cognition and cause subjects to simply choose with a fixed probability and ignore the amounts and delays. This would manifest as a reduced relationship between *k*_*eq*_ and *pSS*. We split the choice set into two halves according to the median *k*_*eq*_ and examined *pSS* for each half, Figure S3. We found a main effect of *k*_*eq*_ on choice in the high magnitude condition, *F*(1,26) = 5.13, *p* = .03, and a trend towards a main effect of *k*_*eq*_ in control, *F*(1,26) = 3.84, *p* = .06. This indicates that subjects are sensitive to the values and delays of the choice. Note that these effects are weaker than our analysis of *k* because this analysis ignores individual differences in discount rate. We next examined whether rTMS influenced the overall probability of choosing the smaller-sooner reward. Relative to sham, we find a main effect of stimulation in the high magnitude condition, F(1,26) = 50, *p* < .001, and a trend towards a main effect of stimulation in control, *F*(1,26) = 3.92, *p* = .06, indicating that subjects are less likely to choose smaller-sooner with stimulation. In both conditions, the interaction between stimulation and *k*_*eq*_ is nonsignificant, both *p* > .2. That is, rTMS does not influence the (estimated linear) relationship between *k*_*eq*_ and choice. In light of these results, we conclude that our overall effects are best interpreted as rTMS having a large effect on choices selectively in high magnitude decisions.

Finally, because previous reports have found an effect of hemisphere of rTMS on discounting ^10^, we also examined whether hemisphere interacted with any of these effects. We found that hemisphere of stimulation had no effect on discounting, no effect on the magnitude effect, no effect on the influence of rTMS on discounting, and no effect on the interaction between rTMS and context, with respect to either control condition (all *p* > .1), Table 1.

**Table 1.** Results of ANOVAs assessing the magnitude effect relative to two different control conditions (Sham and Pre-Experiment). Both ANOVAs show significant effects of magnitude, a significant effect of TMS, and a significant interaction between the magnitude effect and TMS. *H* refers to generalized eta squared.

|  | EFFECT | DF | F | H | P |
| --- | --- | --- | --- | --- | --- |
| SHAM | hemisphere | 1, 25 | 0.35 | 0.005 | 0.56 |
|  | magnitude (mag) | 1, 25 | 43.8 | 0.27 | <.001 |
|  | hemi:mag | 1, 25 | 0.09 | 0 | 0.77 |
|  | tms | 1, 25 | 9.57 | 0.11 | 0.005 |
|  | hemi:tms | 1, 25 | 0.04 | 0 | 0.85 |
|  | mag:tms | 1, 25 | 49.3 | 0.15 | <.001 |
|  | hemi:mag:tms | 1, 25 | 0.71 | 0.003 | 0.41 |
| PRE-EXPERIMENT | hemisphere | 1, 25 | 0.02 | 0 | 0.89 |
|  | magnitude (mag) | 1, 25 | 172 | 0.39 | <.001 |
|  | hemi:mag | 1, 25 | 0.08 | 0 | 0.78 |
|  | tms | 1, 25 | 15.5 | 0.19 | <.001 |
|  | hemi:tms | 1, 25 | 0.98 | 0.01 | 0.33 |
|  | mag:tms | 1, 25 | 90.4 | 0.27 | <.001 |
|  | hemi:mag:tms | 1, 25 | 2.71 | 0.01 | 0.11 |

### 2.2 Hidden Zero Effect

We next turned to the effect of rTMS on the hidden zero effect, which refers to the finding that people are more patient when zero-value rewards are stated explicitly, e.g. $20 now and $0 in two weeks. Previous work has suggested that the hidden zero effect arises due to a difference in reward processing in the striatum between framing conditions^21^. Additionally, this study found no difference in dlPFC activation between hidden and explicit zero framing. We therefore predicted that rTMS would have no impact on the hidden zero effect.

We analyzed data in the same manner as for the magnitude effect. We first tested for the presence of a hidden-zero effect and observed an effect of explicit zero framing on discounting with respect to both pre-experiment baseline, *F*(1,25) = 18.8, *p* < .001, □*g*^2^= 0.06, and sham TMS, *F*(1,25) = 7.0, *p* = .01, □*g*^2^= 0.03, Figure 1. Therefore, we replicated the hidden zero effect.

With respect to sham TMS, we found no interaction between TMS and the framing manipulation (*p* > .2), consistent with our hypothesis. Contrary to our hypothesis, we did observe an interaction between TMS and the hidden zero effect with respect to pre-experimental baseline, TMS, *F*(1,25) = 5.1, *p* = .03, □*g*^2^= 0.02, but not in sham, *p* > .2, Figure 2. It is possible that this difference between controls was due to the difference in the baseline size of the hidden zero effect. We conducted post-hoc tests of the hidden-zero effect in our control conditions and found that it was significant in the pre-experiment baseline, *t*(24) = 3.49, *p* < .002, but only marginal in sham TMS, *t*(24) = 1.93, *p* = .064. Therefore, the lack of a significant effect of rTMS on the hidden zero effect with respect to sham TMS could be driven by the fact that the hidden zero effect was weak in sham TMS. Like the magnitude effect, we found no effects of rTMS on discount rates by hemisphere (Table 2). Overall, our results are inconclusive but suggestive of an unanticipated effect of dlPFC-rTMS on the hidden-zero effect.

**Table 2.**
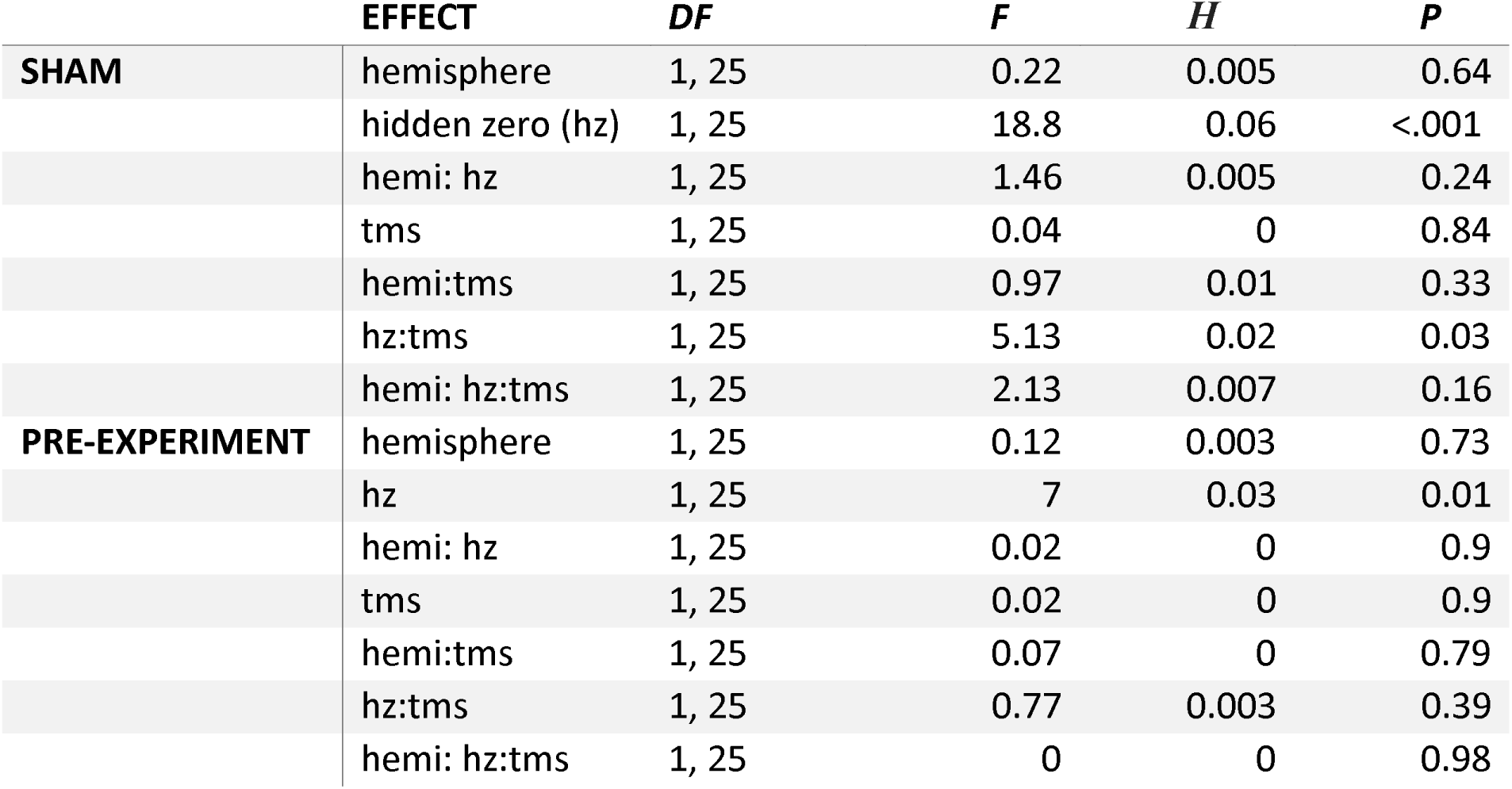
Results of ANOVAs assessing the hidden zero effect relative to two different control conditions (Sham and Pre-Experiment). Both ANOVAs show significant effects of hidden zero framing. However, only the pre-experiment assessment of baseline showed a significant effect of TMS on the hidden zero effect. *H* refers to generalized eta squared.

### 2.3 Direct comparison of effects

Our original hypothesis was that the rTMS would disrupt the magnitude effect but not the hidden zero effect. We found evidence that rTMS disrupted both effects. We next considered whether rTMS had a *larger* effect on the magnitude effect than on the hidden zero effect. However, this hypothesis is difficult to assess in an unbiased manner because the magnitude effect is so much larger than the hidden zero effect in the absence of rTMS. Given the baseline size of the two effects, it is mathematically impossible for rTMS to exert a larger influence on the hidden-zero effect (unless TMS caused the hidden effect to reverse). Note that this bias also effects standard nonparametric tests such as signed ranks. To avoid this, we counted the number of subjects for whom TMS reduced the magnitude effect (26 out of 27) and the number for whom TMS reduced the Hidden Zero effect (18 out of 27) and compared these proportions using a two-sided Fisher’s exact test. We found that the proportions differed, odds ratio = 13.0, *p* = .011, indicating that rTMS reduced the magnitude effect more frequently than it reduced the hidden zero effect. However, the relative weakness of the baseline hidden zero effect makes it impossible to draw strong conclusions from this result.

## 3 Discussion

Cognitive control refers to a set of processes that support goal directed behavior. Control mechanisms likely influence intertemporal choices by maintaining a representation of specific goals that bear on the decision (e.g., I am going on vacation this summer and need to save), building a mental simulation of the hedonic experience of receiving a reward in the future, and guiding attention to the relevant dimensions of the choice^13^. Cognitive control therefore acts as countervailing force against impulsive choice by encouraging a balanced assessment of one’s true preference. However, cognitive control is costly to exert and is adaptively deployed in more important or more difficult decisions^24^,^25^. In the magnitude effect, large valued choices may signal a more important decision and trigger the increased engagement of control processes ^8^. We demonstrated that disruption of dlPFC activity, which has been robustly associated with cognitive control^26^, dramatically reduces the magnitude effect. This result establishes a causal role for the dlPFC in the magnitude effect.

In addition, this result is inconsistent with the predictions of the utility model of the magnitude effect. Because rTMS over dlPFC should influence control processes without directly affecting valuation of rewards^10^, the utility model incorrectly predicts that rTMS should have a similar effect in both high and low magnitude contexts. Our previous work showed that hunger, which is associated with reduced self-control, reduces the magnitude effect by increasing impulsivity for large magnitude choices. In addition, we showed that asking subjects to justify their decisions, which enforces the exertion of self-control, eliminates the magnitude effect by decreasing impulsivity for small magnitude choices^8^. It was possible that these behavioral manipulations effected subjective valuation, and therefore the present rTMS finding provides an important piece of evidence against the utility model. Further evidence against the utility model comes from behavioral assessment of utilities, which have failed to support the utility model’s central tenant that subjects show increasing sensitivity to proportional differences with large magnitudes^27^. Of the other models of the magnitude effect the authors are aware of, only the *memory sampling* model can account for previous behavioral findings as well as the present rTMS result. According to one version of this model, decisions are made by sampling experiences from memory that are similar to the available options and comparing the affective value of those experiences^28-30^. Because larger rewards are generally associated with longer delays (e.g., monthly paychecks), the same time delay will seem comparatively shorter when considered in the context of a large reward and have less of an impact on choice. It is possible that disruption of the dlPFC impairs the ability to search memory for experiences with large, delayed rewards, which are less common than experiences with small rewards. In addition, memory sampling is a broad class of algorithms with different mechanisms for drawing from and evaluating memory samples. To the authors’ knowledge, other memory sampling models, such as MINERVA-DM^31^ or query theory^32^, have not been applied to the magnitude effect. Nonetheless, future work is needed to test the self-control model against these diverse sampling models in paradigms where they make opposing predictions.

In addition to the effect of rTMS on the magnitude effect, we find mixed evidence that rTMS reduces hidden zero effect. Specifically, we found a significant difference of rTMS compared to pre-experimental baseline but not to sham rTMS. However, the hidden-zero effect was weak in the sham condition. There are several possible explanations for this result. First, the hidden zero effect may be sensitive to pre-exposure to the task and therefore may diminish over time. Second, there might be a stronger dependence of the hidden zero effect on prefrontal cortex than previously anticipated. Third, the integration of the additional information (i.e. the explicit zero) may partially depend on functions supported by the dlPFC. Thus, even though the information is explicit, it may be more difficult to attend to and incorporate this information under rTMS. Future work should systematically examine these possibilities.

We did not observe any effects of hemisphere in any of our comparisons. This result contrasts with the finding that rTMS^10^ and current stimulation^33^,^34^ of left, but not right, prefrontal cortex biases decision-making. In spite of these findings, a large literature has established that cognitive control functions are associated with a bilateral fronto-parietal network^35^,^36^. With the exception of response inhibition, which is more strongly associated with right ventrolateral prefrontal cortex^37^, there is little empirical evidence that control mechanisms are lateralized. Our results, although divergent from previous brain stimulation studies of intertemporal choice, are therefore consistent with a broad literature showing bilateral prefrontal involvement in self-control.

A major goal of decision neuroscience is to understand the psychological dimensions and neural processes that underlie decisions between delayed rewards. This in turn could lead to novel strategies for combatting maladaptive decision making. Our finding that the dlPFC is causally involved in the magnitude effect suggests that interventions that increase the function of this region and its associated brain networks could improve economic decision making.

## 4 Method

### 4.1 Participants

Thirty right-handed participants completed this study (11 females; mean age = 26.4 ± 3.8). This sample size was decided a priori and data analysis did not begin until all data were collected. However, we did not conduct a formal power analysis. All participants reported having no history of neurological or psychiatric problems and no females were currently pregnant. All participants gave written informed consent to participate in the study that was approved by and conducted in accordance with the policies of the Stanford University Institutional Review Board. One participant was excluded due to excessive head movement during rTMS, and two of them chose to withdraw during the task, leaving twenty-seven participants for analysis. No participant experienced adverse effects or reported any scalp pain, neck pain, or headaches after the experiment. We randomly assigned participants to receive a train of rTMS to either the left dorsal lateral prefrontal cortex (dlPFC) (left rTMS group; n = 15), or the right dlPFC (right rTMS group; n = 12).

### 4.2 Repetitive Transcranial Magnetic Stimulation (rTMS)

rTMS was administrated to the dlPFC for 15 minutes before participants conducted the main task. Low-frequency (1 Hz) rTMS was delivered with a commercially-available figure-eight coil (70-mm diameter double-circle (Magstim, Winchester MA). The coil was held tangential to the participant’s head, with the handle pointing rostrally. Participants received a single, 15-min, 1-Hz rTMS train (900 pulses) over either the left dlPFC or right dlPFC. We tailored the stimulation strength by stimulating the motor cortex. Once we found the smallest amount of stimulation to make a finger consistently twitch, we set the power to 120% of this motor threshold and gave participants a test stimulation of 1Hz for 30 seconds and asked them if they could tolerate the stimulation. If the participant had a threshold over 83% of maximal TMS power, we ran at 100% power. Sham stimulation was delivered by flipping the coil to stimulate away from the head. This produces the same clicking sound as produced by the real stimulation and significantly lowers the stimulation level. Flipping the coil has commonly been used to redirect the TMS field as a control condition^38^.

For stimulation of the left and right dlPFC, the TMS coil was placed over F4 and F3 using the electroencephalogram 10-20 coordination system, as in previous studies^10^. The rTMS parameters were well within currently recommended guidelines, and stimulation using these parameters results in a suppression of excitability of the targeted cortical region^22^. There is considerable uncertainty about the duration of the rTMS effects. However, effects appear to last for at least as long as half the duration of the stimulation^39^. Therefore, participants were asked to conduct the main tasks immediately after the stimulation in the same laboratory room. Task instruction and practice trials were given before the stimulation. All three conditions took on average less than 8 minutes in pre-test session, and less than 6.5 minutes in session 1 and 2. There was a 20-minute long break between sessions, and all participants received real or sham rTMS to the dlPFC in session 1 and the other type of stimulation to the dlPFC in session 2. We did not record the order of the stimulation protocol for each subject and are unable to examine whether there were effects of TMS that persisted into the Sham session for subjects who were given TMS first. Subjects were blind to the type of stimulation they received.

### 4.3 Choice Tasks and Procedures

The original purpose of the study was to compare the effect of rTMS on two different discounting effects. The first, the magnitude effect, is the phenomenon that people become more patient as the magnitude of all options increase. We refer to this is a *context effect*, because discounting as a function of the proportional difference between rewards changes between low and high magnitude decisions. The second, the hidden zero effect, is the finding that people are more patient when zero valued options are stated among immediate and delayed rewards, e.g. $20 now and $0 in two weeks. We refer to this as a *framing effect*, because the actual decision is the same but the information is presented differently. Based on previous neuroimaging work, we hypothesized the magnitude effect should be more strongly affected by rTMS over dlPFC than the hidden zero effect^8^,^40^. We explain in the Results why differences between the conditions make comparisons difficult to interpret, and we report the results of all experimental conditions.

Three conditions (control, high magnitude, and explicit zero condition) consisted of 20 choices each. All three conditions were scheduled as within-subject design, and the order of conditions was randomized. Subjects made a series of binary intertemporal choices between smaller rewards, *r*_1_, available immediately, and larger rewards, *r*_2_, available after delay *t* (2 weeks or 1 month). In the control and high magnitude condition, the options were simply presented as a form of ‘$*r*_1_ today’ for the sooner option, and ‘$*r*_2_ in *t*’ for the later option. In the explicit-zero condition, the options had “explicit-zero” framing, so the sooner option is expressed as ‘$*r*_1_ today and $0 in *t*’ and the later option is expressed as ‘$0 today and $*r*_*2*_ in *t*.’

In order to construct the choices, we sampled *r*_1_ from a normal distribution *N*(20,5). For each choice set of 20 choices, we constructed *r*_2_ from a set of percent differences (*r*_2_/*r*_1_ – 1): (1%: 3 choices, 5%: 3 choices, 10%: 3 choices, 20%: 3 choices, 30%: 3 choices, 40%: 3 choices, 50%: 2 choices). For example, if the smaller reward was 20, and the percent difference was 20%, the larger reward would be 24. All smaller rewards were available immediately. Half of the larger rewards were available in 14 days and half were available in 28 days. The delays were sampled pseudorandomly, such that each delay had at least one choice with each percent difference in values. Procedures for the hidden zero and the control conditions were identical. For the high magnitude condition, we sampled values from *N*(2000, 500) and the percent differences were reduced by an order of magnitude (.1% - 5%)^8^. This reduction in percent differences is essential because the magnitude effect is so pronounced that fixing proportional differences creates large subjective value differences between conditions (e.g., our average subject is roughly ambivalent between $20 now and $24 in 2 weeks, but overwhelmingly prefers $2,400 in 2 weeks to $2000 now). This design therefore enables us to estimate discount rates for both high and low magnitude conditions and match the difficulty of the choice set between conditions^8^. Because discount rates for hypothetical rewards are a close proxy to those of real rewards^41^, and because discounting effects are consistent between real and hypothetical rewards^41^, we used hypothetical decisions. Earlier options were always presented on the left side of the display and preferences were indicated by button press. For most subjects, choices were presented with decimal values (e.g., $20.45); however, due to a coding error, four subjects have integer smaller-sooner (but not larger-later) conditions. Because this this change affected all conditions for those subjections and we used a within-subjects design, this discrepancy should not affect our results. Choice trials were self-paced with a maximum response time of 15s. After making a selection, subjects viewed a 2s feedback screen indicating that the response was recorded.

Two days before the laboratory experiment, participants completed online choice tasks. This pretest used the same format as the main task, except that there was no time limit. On the date of the in-lab experiment, participants gave written consent for the rTMS treatment, and were instructed about the rTMS procedures.

### 4.4 Analysis

Discount rates, *k*, were estimated for each condition assuming a hyperbolic discount function of the form:

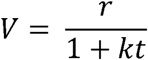

where *V* is the subjective value, *r* is the reward in dollars, and *t* is the time to reward delivery. We estimated a discount rate for each subject individually by assuming a softmax decision function and maximizing the log-likelihood of the observed choices as a function of *k* and the softmax noise (inverse temperature) parameter *α*. This approach utilizes the empirically-validated shape of discount functions to disentangle the effects of choice noise from discount rate on choice. As such, discount rate is a sensitive and specific dependent variable that captures impulsivity. However, we also conducted a similar analysis on the probability of smaller-sooner (*pSS*) choice. This approach conflates discount rate and choice noise, but it is more transparently related to the raw data. The results of this analysis are qualitatively the same as with discount rates. Since the results based on *pSS* are consistent with analyses based on *k*, we report the *pSS* analyses only in Supplemental Information. We fit the models using the conjugate gradient algorithm implemented in Scipy within the Python programming language. In addition, we used Scipy’s basinhopping wrapper with 50 initializations in an attempt to avoid local minima. Discount rates were log-transformed. All tests used type III mixed ANOVAs with Greenhouse-Geisser sphericity correction implemented in R.

### 4.5 Data Availability and Open Practices

All data and analysis code can be accessed at https://github.com/iancballard/tms_discounting

**Author Contributions**
I.B. analyzed the data. I.B. curated the data and wrote the manuscript. B.K. and S.M. conceptualized the design. S.M. provided supervision and S.M. and G.A. provided critical revisions.

## Acknowledgements

This work was supported by NSF grant 1358507 (SMM).

